# Extensive cellular heterogeneity of X inactivation revealed by single-cell allele-specific expression in human fibroblasts

**DOI:** 10.1101/298984

**Authors:** Marco Garieri, Georgios Stamoulis, Emilie Falconnet, Pascale Ribaux, Christelle Borel, Federico Santoni, Stylianos E. Antonarakis

## Abstract

In eutherian mammals, X chromosome inactivation (XCI) provides a dosage compensation mechanism where in each female cell one of the two X chromosomes is randomly silenced. However, some genes on the inactive X chromosome and outside the pseudoautosomal regions escape from XCI and are expressed from both alleles (escapees). Given the relevance of the escapees in biology and medicine, we investigated XCI at an unprecedented single-cell resolution. We combined deep single-cell RNA sequencing with whole genome sequencing to examine allelic specific expression (ASE) in 935 primary fibroblast and 48 lymphoblastoid single cells from five female individuals. In this framework we integrated an original method to identify and exclude doublets of cells. We have identified 55 genes as escapees including 5 novel escapee genes. Moreover, we observed that all genes exhibit a variable propensity to escape XCI in each cell and cell type, and that each cell displays a distinct expression profile of the escapee genes. We devised a novel metric, the Inactivation Score (IS), defined as the mean of the allelic expression profiles of the escapees per cell, and discovered a heterogeneous and continuous degree of cellular XCI with extremes represented by “inactive” cells, i.e., exclusively expressing the escaping genes from the active X chromosome, and “escaping” cells, expressing the escapees from both alleles. Intriguingly we found that *XIST* is the major genetic determinant of IS, and that XIST expression, higher in G0 phase, is negatively correlated with the expression of escapees, inactivated and pseudoautosomal genes. In this study we use single-cell allele specific expression to identify novel escapees in different tissues and provide evidence of an unexpected cellular heterogeneity of XCI driven by a possible regulatory activity of *XIST*.

## INTRODUCTION

In eutherian mammals, X chromosome inactivation (XCI) is a well-described mechanism of dosage compensation for the X chromosome in females (Lyon 1961; Penny et al. 1996; Chow and Heard 2009; Bartolomei and Ferguson-Smith 2011). In female cells, only one X chromosome is transcribed (Xa; X-active), whereas the second X chromosome is silenced (Xi; X-inactive) (Lyon 1961). Marsupials have an imprinted pattern of XCI, and the paternal allele is predominantly inactive (Sharman 1971). In mice, an imprinted form of XCI occurs through early embryonic developmental stages (4-8 cell stage)(Huynh and Lee 2003; Okamoto et al. 2004; Okamoto et al. 2005; Patrat et al. 2009), followed by inner cell mass reactivation and random XCI in epiblast cells (Mak et al. 2004). In humans, the two X chromosomes are active during post-zygotic stages, achieve gene dosage compensation by dampening their expression up to or even after late blastocyst formation, until one of the X chromosomes is randomly inactivated in each cell (Petropoulos et al. 2016). In female somatic cells, random XCI is stable, resulting in a mosaicism for gene expression on the X chromosome, in which an average of 50% of cells express the active paternal X and 50% the active maternal X alleles. Most of the genes on the Xi chromosome are transcriptionally silenced through epigenetic processes initiated by the X Inactivation Center (XIC) and spread along the Xi chromosome during early embryogenesis (Lyon 1961). The XIC encodes several genes, including *XIST*, a long non-coding RNA (ncRNA) essential for initiating and completing XCI (Brown et al. 1991; Ballabio and Willard 1992; Brown et al. 1992). *XIST* RNA molecules mediate the establishment and maintenance of XCI in subsequent cycles of mitotic division by coating the Xi chromosome and recruiting Polycomb Repressive Complex 2 with repressive chromatin modifiers (Lee and Bartolomei 2013). It has been shown that the coating of the Xi is regulated by the interaction between *XIST* ncRNA and the Lamin B receptor (LBR)(Chen et al. 2016a). This interaction is needed for the recruitment of Xi to the nuclear lamina and the subsequent spread of *XIST* ncRNA to actively transcribed regions (Chen et al. 2016a).

However, not all X-linked genes are inactivated. In females, genes that escape from XCI (escapees) represent 15-25% of the X-linked genes, and a further 10% of escapees differ between individuals and cell types (Carrel and Willard 2005; Prothero et al. 2009; Yang et al. 2010; Cotton et al. 2013; Crowley et al. 2015). Such genes have been associated to sex-specific traits and to clinical abnormalities in patients with X chromosome aneuploidy, such as Turner and Klinefelter Syndromes (Berletch et al. 2011). Pathogenic variants in escapees also contribute to various disease phenotypes in women carriers, including Kabuki syndrome (KABUK1 [MIM 147920])(Lederer et al. 2012; Miyake et al. 2013), intellectual disabilities(van Haaften et al. 2009; Grasso et al. 2012; Jones et al. 2012; Gropman and Samango-Sprouse 2013; Zhang et al. 2013; Dunford et al. 2016). Genes escaping XCI have been previously identified by whole tissue studies using different approaches, such as X-linked gene expression comparisons between males and females (Yasukochi et al. 2010), detecting allelic imbalance in clonal lymphoblast and fibroblast cell lines (Cotton et al. 2013),identifying inactivated and active transcription start sites by methylation profiles (Cotton et al. 2015)and among female individuals with X chromosome aneuploidies (Sudbrak et al. 2001)

The ability to capture single cells and to study their allele-specific expression (ASE) (Borel et al. 2015) provides the opportunity to explore XCI patterns at the single-cell level and to identify escapee genes. Recent studies on mouse single cells demonstrated the robust nature of this technology to monitor the dynamics of XCI through differentiation (Chen et al. 2016b), mouse preimplantation female embryos (Borensztein et al. 2017) and in clonal somatic cells (Reinius et al. 2016). Recently, (Tukiainen et al. 2017) performed an across-tissue study of X inactivation and partially validated their observation performing shallow sequencing (1Mio reads x cell) on 940 single cells from lymphoblasts and dendritic cells. Here, using RNA-Seq at high sequencing depth (40Mio reads per cell), we studied the X-linked ASE in 983 isolated, unsynchronized single fibroblast and lymphoblast cells and established the degree of XCI after the removal of potential confounding effects. One of the caveats of allele expression quantification in single cells is represented by the allele dropout, which randomly affects the detection of one of the two alleles of poorly expressed genes (Stegle et al. 2015). However, in this context, the allelic dropout will not induce false positive escapees. Moreover given the number of cells analyzed in our study, the probability to consistently miss the capture of the expressed allele from the X-inactive chromosome of a true escapee in all the cells is extremely low. In this study we identified 55 escapee genes in at least one individual, out of which 5 were novel. A subset of 22 genes was detected as escapee in at least two individuals (robust set), including 3 novel escapee genes. Through the analysis of the expression profile of the genes in the robust escapee set, we further investigated their propensity to escape XCI in each cell. We discovered that the transcriptional activity of the escapee genes is highly variable and significantly associated with the cellular *XIST* transcript abundance.

## RESULTS

### Identification and elimination of confounding doublets

Genes located on the X chromosome of female cells express one allele from the randomly active chromosome, while escaping genes express both alleles (Fialkow 1970). Multiple cells (i.e., doublets) resulting from the simultaneous capture of two or more cells expressing discordant haplotypes obviously complicate the detection of escapee genes and potentially increase the number of false positives. A recent study has explored the limits of current fluidic technology to conduct single-cell RNAseq, demonstrating that a substantial fraction of the bona fide isolated single cells are doublets (Macosko et al. 2015). After removing 32 cells because of low mapping quality (Supplemental Figure S1), in order to eliminate all doublets with discordant haplotypes, we conducted a pairwise correlation analysis of X-linked ASE among all the cells. We performed hierarchical clustering analysis and obtained three distinct clusters of cells (Figure 1B and Supplemental Figure S2, Methods). As expected, two clusters were populated by cells with one mutually inactivated X chromosome. The third cluster included cells with a biallelic expression pattern for all the X chromosome genes, revealing the presence of doublets. Considering all 5 individuals (n = 983 cells), we identified a total of 82 doublets (~8% of the total, consistent with Fluidigm manufacturer’s expectations), which were excluded from further analysis (Figure 1B, Supplemental Figure S2 and Supplemental Table S1). Doublets expressing concordant haplotypes cannot be detected with this our *in silico* approach; however, they do not inflate the number of false positive escapees and, consequently, they do not have an impact on escapee gene discovery and XCI analysis.

**Figure 1:**
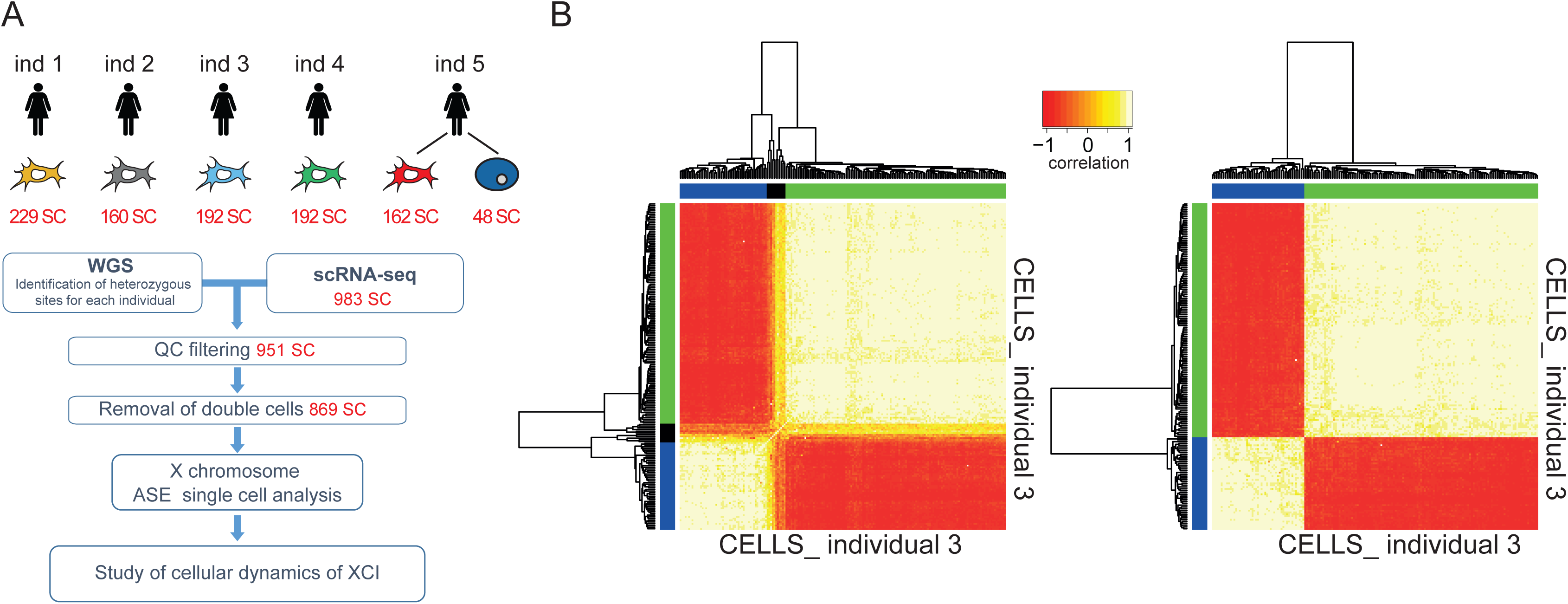
Identification and elimination of confounding doublets. (A) Flowchart of the study. Whole genome sequencing for each individual identified the respective heterozygous sites. Single-cell RNA-seq provided RNA abundance for each single cell. After QC and removal of the doublets, our framework calculated AR for the X-linked genes to enable escapees identification and study the cellular XCI. (B) Heat map of unsupervised hierarchical clustering using cell-cell pairwise Pearson correlation coefficients of AR from single cells from individual 3 before (left) and after (right), excluding the identified doublets. The heat map separated the cells expressing one haplotype (blue and green cluster) from cells expressing two haplotypes (black cluster, doublets). Pearson correlations range from -1 (red) to 1 (white).

### Identification of escapee genes

To estimate the allelic ratio (AR) for each gene in each cell, we calculated the ratio of the number of reads supporting the cell-specific expressed haplotypes over the total number of reads covering all single nucleotide variants (SNV) of a gene (See Methods). Fully inactivated genes displayed an AR equal to 1. In the relaxed discovery set of escapee genes, putative escapees were defined as having an AR =< 0.95 in at least one individual. The rational and the choice of this threshold is explained in the methods. The gene is considered as inactivated (i.e. exclusively expressed from the active chromosome) otherwise. As a proof of principle of the reliability of our approach we first confirmed that *XIST* is expressed exclusively from the inactivated allele by analyzing its AR in all cells from individuals 3 and 4 (i.e monozygotic twin samples), for which we were able to phase the haplotypes from parental genotyping (Supplemental Figure S3). As an additional control we examined the allele expression profile of genes in PAR1 and PAR2 regions. As expected, all of these genes exhibited a balanced ASE across the two X chromosomes except *VAMP7* in PAR2 where very few cells displayed an expression from the inactive chromosome (Figure 2, Supplementary Table S2).

**Figure 2:**
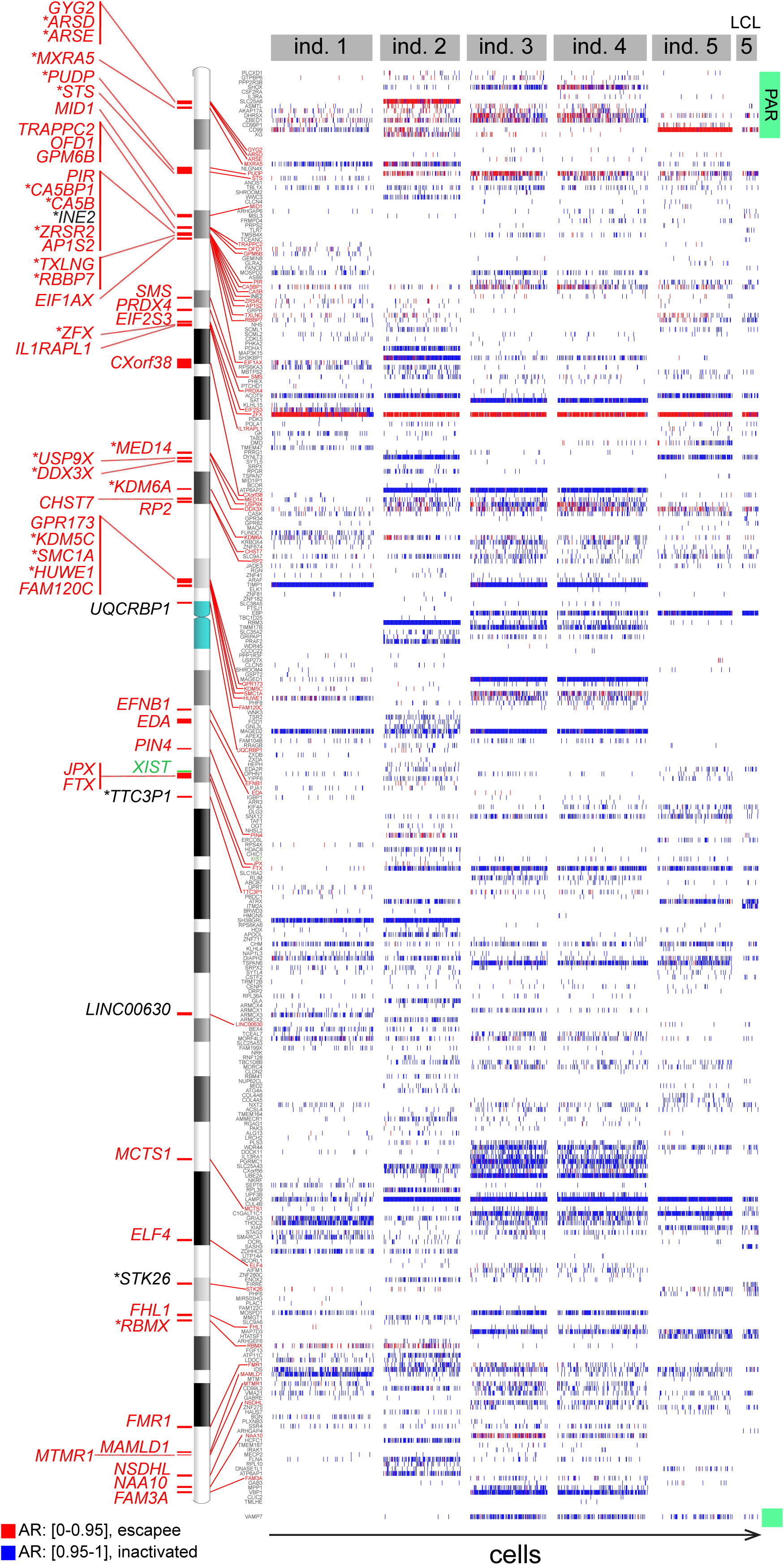
Single cell Allelic Ratio profiles with respect to the active haplotype for genes on female X chromosome. For each individual, the Allelic Ratio for each gene (fibroblasts or lymphoblasts) is reported for each cell along the x-axis (rectangles with AR ≥ 0.95 (blue) an AR ≤ 0.95 (red)) according to the genomic location of genes in the human X chromosome (y-axis). 55 identified escapee genes in at least one individual are annotated on the left of the X chromosome ideogram. Known escapees are shown in red; novel escapees in black; escapees from the robust set with an asterisk. *XIST* is shown in green. PAR, pseudoautosomal regions; LCL, lymphoblastoid cells.

As expected, the majority of chrX genes showed an inactivated status in all the cells (Figure 2 and Supplemental Table S2). From a total of 296 genes interrogated in at least one individual, we identified the relaxed set of 55 escapees (18.5%): 50 of them previously described to escape XCI in at least one study, and 5 novel escapees *(INE2* (antisense gene), *STK26, UQCRBP1, LINC00630* and *TTC3P1)* (Figure 2, S3). Out of 203 genes with AR information from at least 2 individuals, we classified as robust escapees 22 genes (10.9%) which exhibit an escapee status in at least two individuals including 3 novel genes *(INE2, STK26* and *TTC3P1)*. As expected, the power to detect a gene escaping XCI is linearly related with the respective expression level (Supplemental Figure S4). The number of overlapping escapees among all individuals is shown in a Venn diagram (Supplemental Figure S5). Results from both relaxed and robust sets are consistent with the current understanding of XCI, in which is predicted that 10% to 20% of X chromosome genes escape XCI (Carrel and Willard 2005). In addition, we analyzed 48 single cells from a lymphoblastoid cell line derived from the individual 5 to investigate the escapee concordance with the fibroblasts. After quality control and doublets removal, we were able to classify 9 genes as escapee in lymphoblastoid cells with 5 of them being known escapees *(DDX3X, KDM6A, MSL3, PUDP, ZFX)*, and 4 novel escapee genes *(IDS, SLC9A7, STAG2, STK26)*. We observed that, though expressed and having an informative heterozygous site, the *MSL3, IDS, SLC9A7*, and *STAG2* genes were not classified as escapees in fibroblasts. Conversely several genes detected as escapees in fibroblasts were not escapees in lymphoblast cells. This could be partially ascribed to the previously observed tissue specificity of XCI (Deng et al. 2014b; Tukiainen et al. 2017) and to the recently discovered peculiar maintenance of XCI in lymphoblast cells **(Wang et al. 2016)**. Additionally, the relatively small number of lymphoblastoid single cells along with the absence of informative heterozygous sites for some genes in other individuals, can affect our ability to detect escapees(Santoni et al. 2017) and to assess and establish their ASE. However the *STK26* escapee status was clearly confirmed in both fibroblast and lymphoblastoid single cells.

### Heterogeneity of escaping XCI

Among five individuals, the 22 robust escapee genes exhibited a heterogeneous AR profile, being inactivated in some cells and escaping XCI in others (Figure 3). Specifically, we calculated a cellular escaping ratio per escapee gene as the proportion of cells escaping XCI with respect to the total number of cells expressing the gene. Some genes displayed a stable cellular escaping ratio among all the individuals, while others were more variable. For example, *CA5BP1* showed consistent cellular escaping ratios ranging from 37% to 52%; *ZFX*, a known constitutive escapee (Schneider-Gadicke et al. 1989), presented with cellular escaping ratios ranging from 85% to 100%, while *DDX3X* had a broader range going from 29% to 61%. Overall, escapees had different cellular escaping ratios, thereby suggesting that each escapee gene is independently regulated.

**Figure 3:**
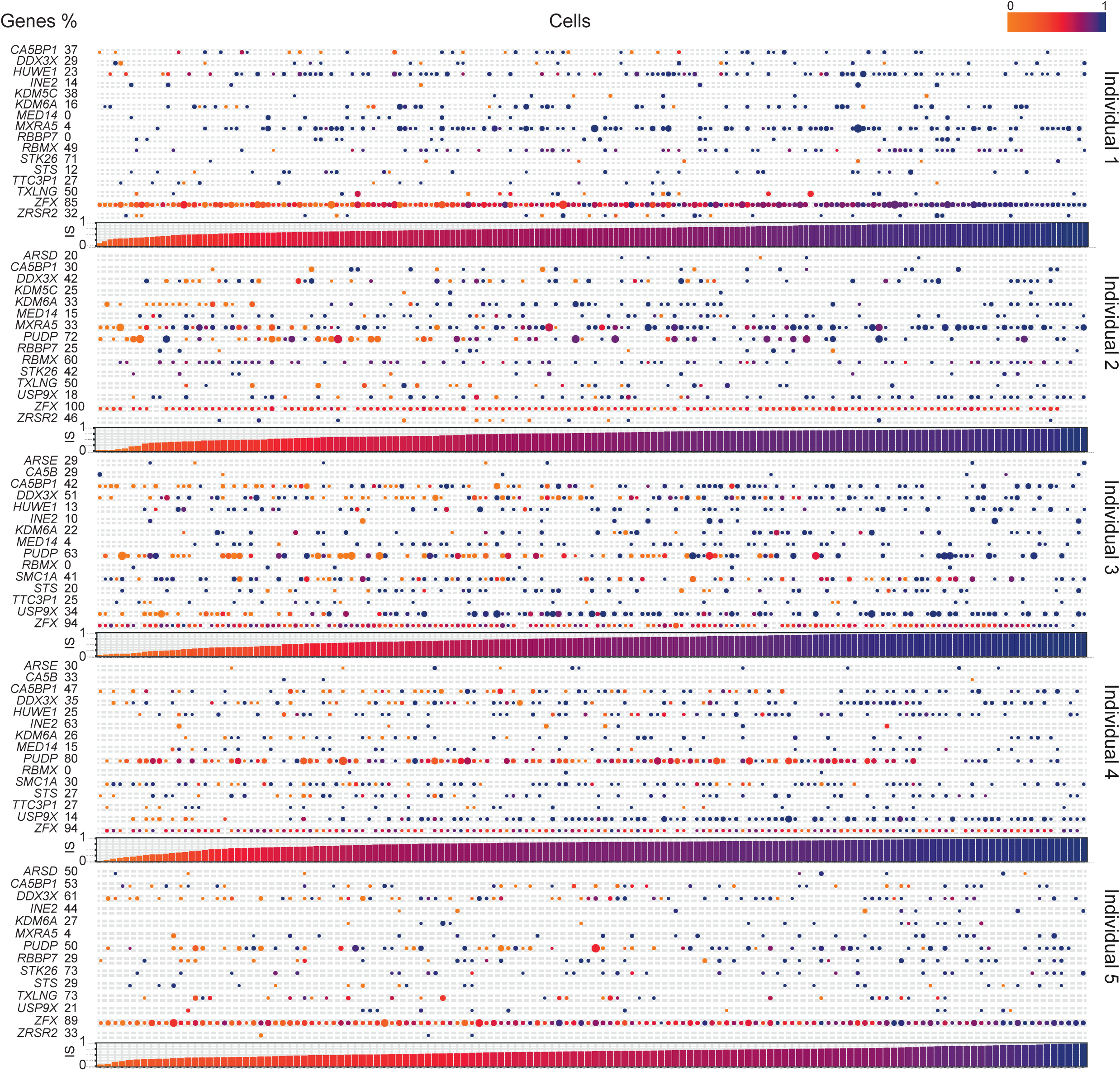
Single cell ASE profile of 22 robust escapee genes in human female fibroblasts. Composite figure of individual allelic ratios per gene per cell (Top of the panel). Allelic Ratio profile of robust escapee genes (listed the rows) with a detectable expression in single cells (ordered along the columns) is shown. Every dot represents the allelic ratio of the respective gene in a cell. AR ranges from 0 (light orange) to 1 (dark blue). The size of the dot is proportional to the respective number of reads detected per cell. (%) is the percentage of cells where the respective gene is escaping XCI. (Bottom of the panel). Bar plot representing the Inactivation Score (see text for details) per cell. IS ranges from 0 (light orange) to 1 (dark blue).

The observed cellular pattern of XCI of the escapees (Figure 3) suggests a variable cellular ability of expressing genes from the inactivated allele. To investigate this hypothesis, we calculated the Inactivation Score (IS) for each cell defined as the mean AR of the escapee genes detected per cell (we considered only cells expressing at least two escapees). For each individual, we ordered the cells according to the respective IS and discovered that the capacity to express from the inactive chromosome is a continuous variable (Figure 3, Figure 4A). This suggested a cellular stratification that reflect the propensity of each cell to escape XCI, as confirmed by the proportion of escapee genes per cell (Figure 4B). As expected, the Inactivation Score is strongly negatively correlated with expression from the inactive X chromosome (Supplemental Figure S6). Notably, in all five individuals, we observed two special groups of cells: one in which all the detectable escapees behaved as inactivated and another where all detectable escapees expressed both X alleles (Figure 4B). These two extreme cell populations represent on average 15% of the total number of aggregate cells among the individuals (individual 1: 20%, individual 2: 14%, individual 3: 8%, individual 4: 7%, individual 5: 26%). As a control, we calculated the Inactivation Score of the remaining inactivated genes per cell (all close to 1, as expected. Figure 4A, right panel). Overall, these results demonstrate that XCI is a complex intra- and inter-individual heterogeneous process, and the ability to escape from X inactivation varies from gene to gene, from cell to cell and also among individuals. The evidence of similar cell stratification in all individuals suggests the existence of a general regulatory mechanism that controls the propensity of a cell to express genes from the inactivated allele.

**Figure 4:**
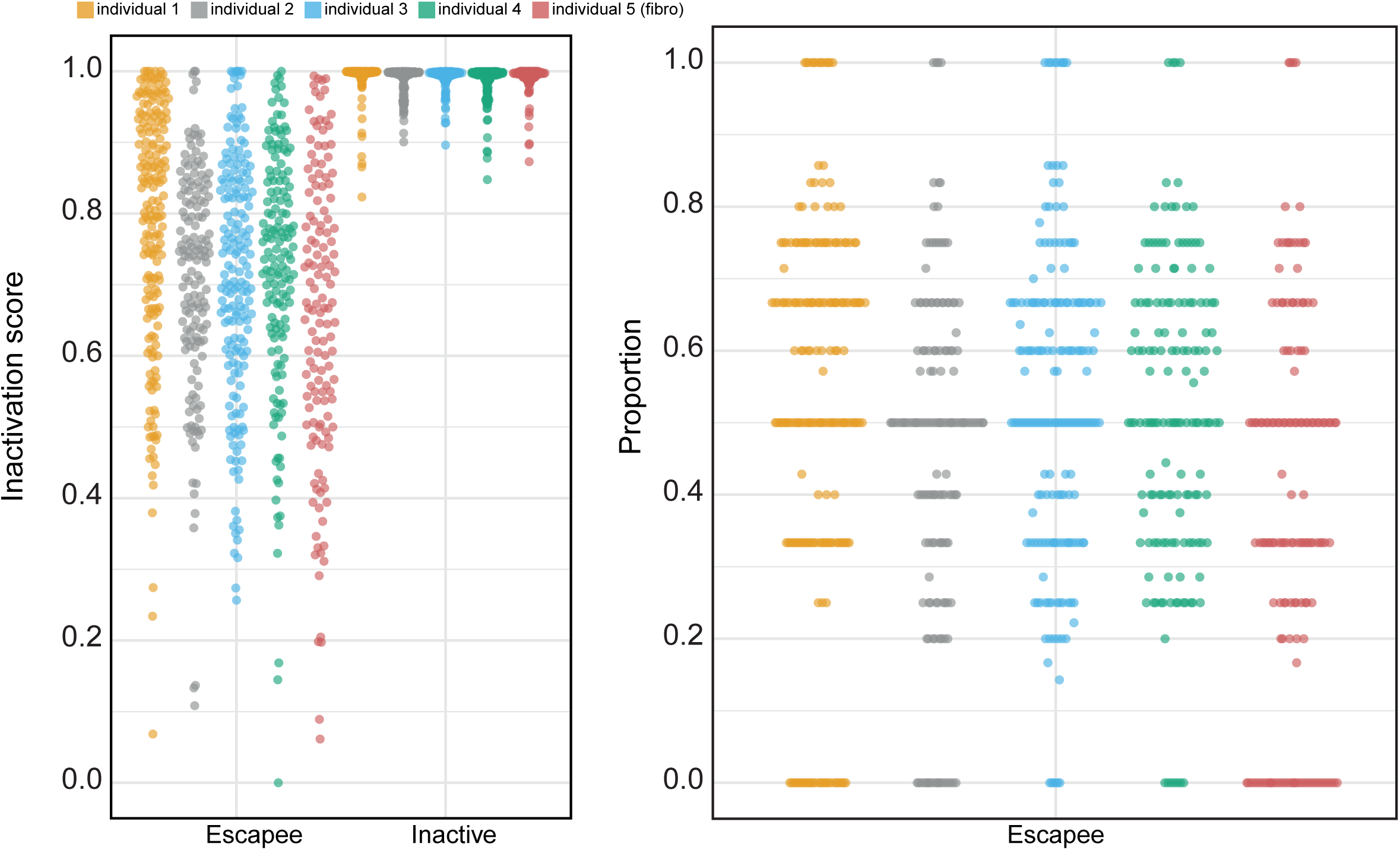
Cellular propensity to escape XCI. (A) Cells ranked by the Inactivation Score calculated on all escapees in the robust set (left - Escapee) and on all inactive genes (right - Inactive) per individual (color coded - see legend). Each dot represents a fibroblast cell. (B) Y-axis-cells ordered according to the percentage of escapees displaying an inactive status (AR ≥\ 0.95, with respect to all detectable escapees) per individual (x-axis). Each dot represents a fibroblast cell.

### Potential drivers of cellular XCI heterogeneity

We hypothesized that the cellular XCI heterogeneity may be associated with the level of expression of genes on the X chromosome. To explore this hypothesis, we correlated the X-linked gene expression (RPKM > 1) with the Inactivation Score. After FDR correction for multiple testing, genes were ranked based on the adjusted p-value (Figure 5A). Notably the only gene positively and significantly correlated with IS was *XIST* (nominal p-value = 3.2 x 10^-5^; adj. p-value = 3.0 x 10^-3^). *XIST* is a well-known non-coding RNA that regulates the establishment and the maintenance of XCI **(Brown et al. 1991)**. Other significantly albeit negatively associated genes were *EIF2S3*, *CD99, NDUFA1, RPL10* and *BCAP31*. We further investigated the correlation of *XIST* expression with inactivated genes and all escapee genes (relaxed and robust sets) and genes located in PAR regions. Notably we observed a significant trend of negative correlations between *XIST* expression and the majority of the genes for all four categories (H0: mean of distribution of correlation coefficients = 0; inactivated genes p = 2.06 x 10^-83^; PAR genes p = 6.49 x 10^-4^; Relaxed escapee p= 1.01 x 10^-^; Robust escapees p= 4.19 x 10^-4^, Mann-Whitney two-tailed test, Figure 5B). This suggests a general repressive *XIST* regulatory effect in all classes of X-linked genes. To further investigate whether this effect was cell cycle dependent, we used Cyclone (Scialdone et al. 2015) to assign each cell to a specific cell cycle phase according to the expression of the appropriate gene markers (from CycleBase (Santos et al. 2015)). Cells not expressing *MKI67* have been previously reported to be in G0 (Sobecki et al. 2017). A statistically significant difference between the populations of single cells in G0 and G1 was observed regarding *XIST* expression (Figure 6). In agreement with our previous observations, the Inactivation Score was also significantly higher in G0 than in G1 cells on average (Figure 6). These results together support the hypothesis that *XIST* ncRNA tends to be more expressed in the resting G0 phase than in G1 and, consequently, the expression of the escapees from the inactive chromosome is reduced in this cell cycle stage. We could not identify any cell-cycle driven effect for the remaining cell cycle stages, likely due to a limited statistical power (small number of single cells classified in these cell cycle stages, Figure 6).

**Figure 5:**
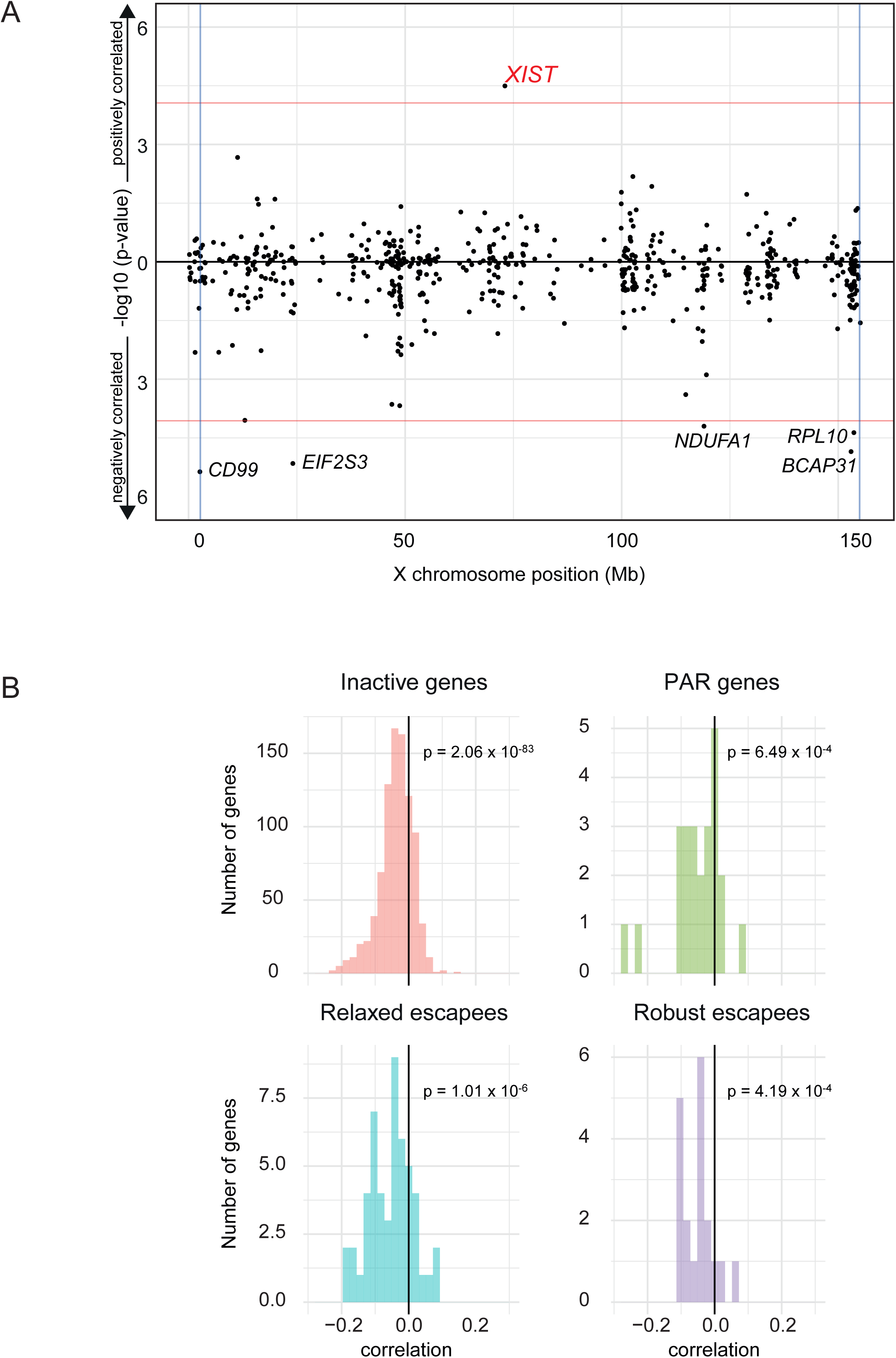
XIST association and correlation with IS and its repressive role for X chromosome genes. (A) Manhattan-like plot for Pearson correlation of X-linked genes expression with Inactivation Score. The plot represents -log_10_ p-value against X chromosome position. Vertical lines represent PAR region limits. Red horizontal lines represent the significance threshold for positively and negatively correlated genes, respectively. (B) Distributions of correlations of expression of XIST with inactivated genes (upper left), PAR located genes (upper right), escapees in robust set (lower left), escapees in relaxed set (lower right). P-values calculated with Mann-Whitney test (see text for details).

**Figure 6:**
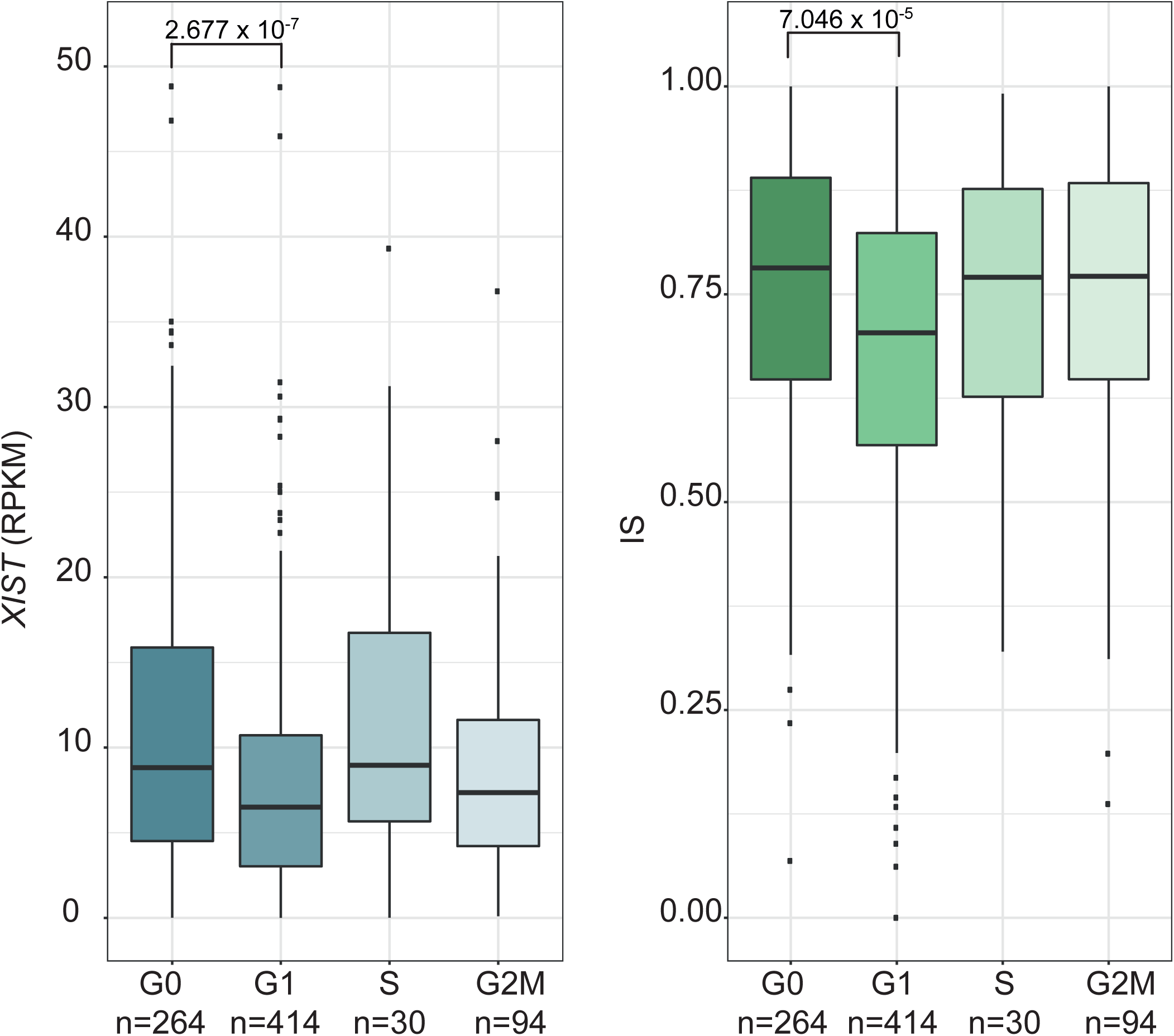
XIST expression and IS dependency on cell cycle. Distribution of *XIST* expression (left) and of Inactivation Scores per single cells (right) according to cell cycle phases (n=number of cells per stage). P-values calculated with Mann-Whitney test (see text for details)

## DISCUSSION

Our study, using human single-cell RNAseq datasets, points to a pervasive heterogeneity in escaping XCI. We have shown that escapees have a different allelic expression profile in single fibroblasts from the same individual. More than 50% of the escapees had the tendency to be mainly expressed from Xa (Figure 4), while *ZFX* and *PUDP* exhibited an overall biallelic expression in more than 70% of the cells. We also observed that some escapees exhibited a relatively stable cellular escaping ratio (proportion of cells in which the gene is escaping), i.e., *ZFX* and *CA5BP1*, where others, such as *DDX3X*, showed broader variability (Figure 3). This finding explains, from a single-cell perspective, the previous observations of heterogeneous gene expression from Xi in cell lines derived from different individuals (Carrel and Willard 1999) and tissues (Cotton et al. 2013; Berletch et al. 2015). A more recent study revealed the contribution of 6 escapee genes *(TRX, CNKSR2, DDX3X, KDM5C, KDM6A, MAGEC3)* to cancer sex bias (Dunford et al. 2016). Here, we confirmed the escapee status for *DDX3X, KDM5C* and *KDM6A*. We observed that the inter-individual *DDX3X* inactivation profile is highly variable and thereby could potentially be associated with differences in cancer predisposition occurring among female individuals.

Among the genes that we detected as escapees in this study, *HUWE1* encodes for an E3 ubiquitin ligase and microduplications of this gene have been found in individuals affected by Turner type intellectual disability (MRX17 [MIM 300706])(Froyen et al. 2008; Froyen et al. 2012). Moreover a recent study suggested the critical role of *HUWE1* as a colonic tumor suppressor gene (Myant et al. 2017). *RBMX* encodes an RNA binding protein which is involved in pre- and post-transcriptional regulation and alternative splicing of many premature transcripts. Pathogenic variants in *RBMX* have been associated with X-linked intellectual disability (Shashi et al. 2000; Shashi et al. 2015). *STK26* is a member of germinal center kinase subfamily and encodes a serine/threonine kinase expressed highly in human lymphocytes and thymus (Ceccarelli et al. 2011; Jiao et al. 2015). Expression of *STK26* (alternatively named as *MST4*) was recently shown to be significantly lower in patients affected by Graves’ disease, an autoimmune disorder characterized by abnormal thyroid function (Guo et al. 2017). It has also been shown that patients affected with Turner Syndrome (XO) exhibit a high prevalence of hypothyroidism (El-Mansoury et al. 2005). We speculate that this phenotype may be related to the reduced expression of the single copy of *STK26*.

Mutations and deletions of the escapee genes identified in the relaxed set such as *EDA, FHL1, IL1RAPL1*, and *FMR1* have been associated with different pathological conditions. More specifically pathogenic variants of *EDA*, a gene of the tumor necrosis factor family, have been associated with X-linked dominant tooth agenesis and X-linked ectodermal dysplasia (Kere et al. 1996) (Yotsumoto et al. 1998) (Visinoni et al. 2003) (Tao et al. 2006) (Tarpey et al. 2007). *FHL1* encodes a member of the four-and-a-half-LIM-only protein family and contains two highly conserved zinc finger domains. Pathogenic variants in *FHL1* have been associated with X-linked dominant reduced body myopathy with both affected male and female individuals (Liewluck et al. 2007) (Schessl et al. 2008; Shalaby et al. 2009), Emery-Dreifuss muscular dystrophy (Gueneau et al. 2009), and scapuloperoneal myopathy (Quinzii et al. 2008). Pathogenic variants of *IL1RAPL1*, a member of interleukin 1 receptor family, have been linked to X-linked mental retardation (Tabolacci et al. 2006; Nawara et al. 2008). A trinucleotide repeat expansion (CGG) in the 5’ UTR of *FMR1* is the cause of Fragile X syndrome (Kremer et al. 1991). We hypothesize that the phenotypic variability of these diseases may be partially explained by variable penetrance and variable expressivity issues in these gene-disease associations due to the observed variable escapee status of their respective causative genes in the relevant tissues.

It has been recently observed in a mouse study that some genes escapee XCI in a fraction of cells only (Reinius (Reinius et al. 2016)et al. 2016). We confirmed and extended this observation with the finding that XCI heterogeneity extends from genes to single cells. Using the Inactivation Score as a metric for cellular propensity to escape XCI, we indeed showed that cells from the same individual have a variable propensity to transcribe the escapee genes from Xi. According to the IS, those cells stratify into three classes: cells where all escapees are expressed from the inactive allele, cells where all escapees are inactivated, and cells with an intermediate profile (Figure 4B). We have shown that the observed cellular heterogeneity is driven by the expression profile of *XIST* in single cells which is in turn strongly and positively correlated with the Inactivation Score. We found five genes negatively associated with IS. *EIF2S3* encodes the gamma subunit of the translation initiation factor 2 which is involved in protein synthesis initiation **(Moortgat et al. 2016)**. This gene has been described to escape X inactivation while hemizygous mutations have been associated with X mental retardation Borck type (Ehrmann et al. 1998) (Moortgat et al. 2016). *CD99* is located in the PAR1 region and is involved in leukocyte migration, and T-cell adhesion (Bernard et al. 1997) (Bernard et al. 2000). Additionally it has been characterized as a potential oncosuppressor in osteosarcoma(Manara et al. 2006). *NDUFA1* (NADH dehydrogenase (ubiquinone) 1 alpha subcomplex, 1) encodes a component of the mitochondrial respiratory chain; hemizygous mutations have been associated with mitochondrial complex I deficiency (Fernandez-Moreira et al. 2007) (Ambros et al. 1997; Potluri et al. 2009). *RPL10* is involved in protein synthesis and is an essential component of large 60s ribosomal subunit (Gong et al. 2009). Hemizygous *RPL10* mutations have been associated with syndromic X-linked mental retardation (Thevenon et al. 2015). *BCAP31* is involved in the export of secreted proteins and hemizygous loss of function mutations have been identified in individuals affected with deafness and severe dystonia **(Cacciagli et al. 2013)**.

Additionally, we demonstrated that *XIST* expression is negatively correlated with the overall expression of all genes in chrX including pseudoautosomal and escapee genes, thereby indicating a general repressive regulatory activity of *XIST* transcripts. Inspired by a previous report describing the localisation of XIST during the cell cycle (Jonkers et al. 2008), we explored the hypothesis that *XIST* expression and IS are related to the cell cycle phases of individual cells. Our data suggest that *XIST* is more expressed during the G0 than the G1 phase and single cells in the G0 phase have an increased IS, indicating less propensity to express escapees from Xi. Overall, these results revealed a considerable degree of heterogeneity of XCI in single cells and enhance our understanding of X inactivation. The framework presented in this study can be applied on an increased number of cells extracted from various tissues from a large cohort of individuals. Such investigations will enable us to define the extent of cellular XCI heterogeneity and clarify its possible implication in the phenotypic variability of X-linked single gene disorders, whole X chromosome aneuploidies and to the observed female sex bias in cancer.

## MATERIAL AND METHODS

### Ethical statement

The study was approved by the ethics committee of the University Hospitals of Geneva, and written informed consent was obtained from both parents of each individual prior to the study.

### Samples

We established 6 different cell lines from 5 female individuals: 5 primary fibroblast cell lines and one lymphoblastoid cell line (Table S1). We captured 935 single-cell fibroblasts and 48 lymphoblastoid single cells. Lymphoblastoid cells obtained from one of the five female individuals (Figure 1A, Table S1)(Borel et al. 2015; Santoni et al. 2017).

### Cell growth

Cells were cultured in DMEM GlutaMAX™ (Life Technologies) supplemented with 10% fetal bovine serum (Life Technologies) and 1% penicillin/streptomycin/fungizone mix (Amimed, BioConcept) at 37°C in a 5% CO_2_ atmosphere as described (Borel et al. 2015).

### Whole Genome Sequencing

Genomic DNA was extracted for all five individuals using a QIAamp DNA Blood Mini Kit (Qiagen) and fragmented by Covaris to peak sizes of 300-400 bp. Libraries were prepared with TruSeq DNA kit (Illumina) using 1 μg of gDNA and sequenced on an Illumina HiSeq 2000 machine with 2 x 100-bp as previously described(Borel et al. 2015). All experiments were performed using the manufacturer’s protocols. For each individual, raw whole genome DNA sequences were analyzed using an in-house pipeline that utilizes published algorithms in a sequential manner (BWA) (Li and Durbin 2010) for mapping the reads over the hg19 reference, SAMtools (Li et al. 2009) for detection of heterozygous sites, and (ANNOVAR) for the annotation (Wang et al. 2010).

### Single-cell capture

Single-cell capture was performed using the C1 single-cell auto prep system (Fluidigm) following the manufacturer’s procedure(Borel et al. 2015). The integrated fluidic circuit used for the study is the C1™ Single-Cell mRNA Seq IFC, 17-25 μm with a capacity of 96 chambers. During the capture, all 96 chambers were inspected under an inverted phase contrast microscope; only chambers containing a non-damaged single cell were considered for downstream analysis.

### Single-cell RNA-sequencing

SMARTer Ultra Low RNA kit for Illumina sequencing (version 2, Clontech) was used for the cell lysis and cDNA synthesis. Libraries were prepared with 0.3 ng of pre-amplified cDNA using Nextera XT DNA kit (Illumina) as described (Borel et al. 2015). Libraries were sequenced on an Illumina HiSeq2000 sequencer as 100 bp single-ended reads. RNA sequences were mapped with GEM (Marco-Sola et al. 2012). Uniquely mapping reads were extracted by filtering for mapping quality (MQ>=150). An in-house algorithm was used to quantify RPKM expression using GENCODE v26. Cells with less than 1 million uniquely mapped reads were excluded from further analysis (Supplemental Figure S7).

### Allele-specific expression and classification of escapee genes

For each gene on the X chromosome, the aggregate monoallelic ratio (AR) per cell was calculated by averaging the allelic ratio of the reads covering the respective heterozygous sites (AR = sum of number of reads from the active X allele / total SNV reads; 0≤AR≤1). Since we do not have the availability of parental genotype for all the individuals, we designed an algorithm to estimate the active X allele per site based on the assumption that the active X allele is, on average, more transcribed than the inactive X. We validated this assumption comparing the prediction of our algorithm with the phasing of twins’ X alleles based on parental information (more details in Supplemental Text S1). According to this metric, inactivated genes cluster around AR=1 while known escapees appear as been uniformly distributed between 0.5≤AR≤0.95 (linear phase of the cumulative distribution, Supplemental Figure S7). As support of this observation, AR distribution of autosomal genes clearly indicates AR=0.95 as the threshold separating biallelically expressed genes from monoallelic expressed genes (Supplemental Figure S8). Therefore, we consider a gene as escapee in the relaxed set when the aggregate AR is ≤ 0.95 in at least 1 individual and as escapee in the robust set when the aggregate AR is ≤ 0.95 in at least 2 individuals. To reduce the effect of allele dropout, we only consider for the analysis SNV sites covered by at least 5 reads in at least three cells. To reduce sampling bias effects (Deng et al. 2014a) a gene is included in the analysis if detectable in more than 5 different cells and/or SNVs per sample.

### Haplotype and multiple cells (doublets) detection

For each cell, the expressed haplotype was estimated by calculating the allelic ratio of each heterozygous site in the X chromosome as identified by whole genome sequencing, excluding sites in the PAR regions (PAR1: chrX:60001-2699520, PAR2: chrX:154931044-155260560) and in known escapee genes (see section Annotation of known escapee genes). The estimated haplotype of each cell was compared to all others through pairwise correlation based hierarchical clustering procedure. A comparison of cells expressing concordant and discordant haplotypes results in a correlation near 1 and -1 respectively. Doublets simultaneously expressing both haplotypes cluster around an absolute correlation of 0.5 are identified and excluded from further analysis.

### Annotation of the escapee genes

First, we curated a list of 190 previously observed escapee genes in different cell types and tissues according to the literature (Carrel and Willard 2005) (Johnston et al. 2008; Park et al. 2010; Yasukochi et al. 2010; Sharp et al. 2011) (Cotton et al. 2013; Zhang et al. 2013)(Table S3). Specifically, we investigated the status of 115 known escapee genes with available informative heterozygous sites and being expressed in fibroblast and lymphoblast cell lines (Supplemental Table S3). Second, we have appended the results published in two studies (Balaton et al. 2015; Tukiainen et al. 2017) in Table S2. Genes detected as escapees in our studies, in absence of citation, have been labeled as novel escapee genes. Genes found as escapee in our study and found subject to inactivation in other studies have been labeled variable escapee genes.

### Cell cycle phase assignment

G1, S, and G2M cell cycle stage related gene markers were obtained from CycleBase (Santos et al. 2015) Cells not expressing *MKI67* have been considered to be in G0 (Schonk et al. 1989) The remaining cells were assigned to their respective cell cycle phase according to the expression of CycleBase genes with Cyclone (Scialdone et al. 2015).

## ACKNOWLEDGMENTS

We thank the laboratory of E. T. Dermitzakis for discussions. This work was supported by grants from the Swiss National Science Foundation (SNF-144082), and the European Research Council (ERC-249968) and the ChildCare foundation to S.E.A. The computations were performed at the Vital-IT (http://www.vital-it.ch) Center for high-performance computing of the SIB Swiss Institute of Bioinformatics.

## ACCESSION NUMBERS

Newly generated RNA and DNA sequencing data are deposited in the European Genome-phenome Archive (EGA, https://www.ebi.ac.uk/ega/) for controlled accesses; the study accession number is (EGASxxxx, to be determined).

## Supplemental Data

Supplemental data include eight figures, additional methods and three tables.

